# *Toxoplasma gondii* injected neurons localize to the cortex and striatum and have altered firing

**DOI:** 10.1101/2021.02.18.431839

**Authors:** Oscar A. Mendez, Emiliano Flores Machado, Jing Lu, Anita A. Koshy

## Abstract

*Toxoplasma gondii* is an intracellular parasite that causes a long-term latent infection of neurons. Using a custom MATLAB-based mapping program in combination with a mouse model that allows us to permanently mark neurons injected with parasite proteins, we found that *Toxoplasma*-injected neurons (TINs) are heterogeneously distributed in the brain, primarily localizing to the cortex followed by the striatum. Using immunofluorescence co-localization assays, we determined that cortical TINs are commonly (>50%) excitatory neurons (FoxP2^+^) and that striatal TINs are often (>65%) medium spiny neurons (MSNs) (FoxP2^+^). As MSNs have highly characterized electrophysiology, we used *ex vivo* slices from infected mice to perform single neuron patch-clamping on striatal TINs and neighboring uninfected MSNs (bystander MSNs). These studies demonstrated that TINs have highly abnormal electrophysiology, while the electrophysiology of bystander MSNs was akin to that of MSNs from uninfected mice. Collectively, these data offer new neuroanatomic and electrophysiologic insights into CNS toxoplasmosis.

## Introduction

A select number of highly divergent microbes (e.g. measles virus, *Toxoplasma gondii*) naturally cause infections of neurons within the nervous system. While most of these viruses cause debilitating disease or death, alpha herpes viruses and the eukaryotic parasite *Toxoplasma gondii* primarily establish persistent, relatively quiescent neuronal infections in those with fully intact immune systems (Dubey, 2009). While neuron-alpha herpes virus interactions have been studied for decades— leading to a mechanistic understanding of herpes virus latency in neurons and the development of novel tools for circuit tracing— our understanding of neuron-*Toxoplasma* interactions is just beginning (Ugolini, 1995; Wickersham et al., 2007).

*Toxoplasma gondii* is a ubiquitous intracellular parasite that can infect a wide range of warm-blooded hosts, including birds, rodents, and humans. In most immunocompetent hosts, *Toxoplasma* establishes a long-term, asymptomatic infection in specific organs. In humans and rodents, the central nervous system (CNS) is a major organ of persistence and is the organ most affected by symptomatic disease in immunocompromised patients (Remington & Cavanaugh 1965; Luft & Remington, 1992; Dubey, 2009; Neu et al., 2015). Using a Cre-based mouse model in which host cells injected with *Toxoplasma* protein permanently express a green fluorescent protein (GFP), our group showed that *Toxoplasma* predominantly interacts with neurons *in vivo* (Koshy et al., 2012; Cabral, Tuladhar et al., 2016). These data suggest that *Toxoplasma* persists in neurons, in part, because parasites primarily interact with neurons rather than glia. In addition, we determined that neurons injected with *Toxoplasma* protein outnumber cysts by over 20-fold (Koshy et al., 2012), suggesting that far more neurons interact with *Toxoplasma* than are persistently infected.

Here we continue to harness this Cre-based model to extend our understanding of the neuron-*Toxoplasma* interface. We leverage the high numbers of *Toxoplasma*-injected neurons (TINs) and the expression of GFP, which labels the full neuron, to carry out a systematic neuroanatomic mapping of TINs. These studies revealed that TINs are most commonly found within the cortex followed by the striatum and are rarely found in the cerebellum. Within the cortex and the striatum, we used immunofluorescent assays to determine if *Toxoplasma* preferentially interacts with specific neurons subtypes, finding that TINs co-localize with markers for the most abundant neuron subtypes in a region. Finally, we used single neuron patch-clamping in *ex vivo* slices to compare the electrophysiology of striatal TINs to neighboring “bystander” neurons (neurons within the same slice that were not injected with *Toxoplasma* proteins). These first-of-their kind study for any neurotropic microbe showed that injection with *Toxoplasma* protein— either directly or indirectly— dramatically altered neuron electrophysiology, while the electrophysiology of bystander neurons remained relatively unchanged.

## Results

### *Toxoplasma*-injected neurons are enriched in the cortex, irrespective of infecting *Toxoplasma* strain

Prior work in AIDS patients and rodents suggested that *Toxoplasma* does not evenly distribute across the brain (Post et al., 1983; Lang et al., 1989; Luft & Remington, 1992; Arendt et al., 1999 Porter & Sande 1992; Strittmatter et al., 1992; Berenreiterova et al., 2011; Evans et al., 2014; Neu et al., 2015; Dubey et al., 2016). These studies have conflicting findings as to where *Toxoplasma* lesions or cysts (in rodent studies) are most commonly found, differences that might be driven by the small numbers of patients or rodents analyzed (Berenreiterova et al., 2011; Evans et al., 2014; Dubey et al., 2016; Boillat et al., 2020). Given these conflicting studies, we sought to neuroanatomically map the location of *Toxoplasma*-injected neurons (TINs) using our previously published semi-automated method (Mendez, Potter et al., 2018). This method utilizes the Allen Institute Mouse Brain Atlas for mapping and allows us to quantify and map TINs at a relatively rapid speed. This automation made it feasible to sample multiple sections of brain per mouse across multiple cohorts of mice infected with either of two genetically distinct *Toxoplasma* strains (type II-Prugniaud, type III-CEP) that express a *Toxoplasma*:Cre fusion protein and mCherry (Koshy et al., 2010, Koshy et al 2012). For simplicity, from here forward, we will refer to these strains as II-Cre and III-Cre. We quantified TINs in 16-19 brain sections/mouse across 32 regions identified in the Allen Institute Mouse Brain Atlas. Data points were grouped into 12 major regions. For both II-Cre or III-Cre infected mice, the cortex housed the highest average number of TINs, followed by the striatum (**Figure 1 A, B**). In general, III-Cre infected mice showed a higher overall number of TINs compared to II-Cre infected mice (average of 247.5 III-Cre vs. 216.7 II-Cre). The increase in absolute numbers in III-Cre infected mice was most pronounced in the cortex (**Figure 1 A, B**). Most other brain regions, including the cerebellum, housed relatively few TINs (**Figure 1 A, B**).

**Figure 1.**
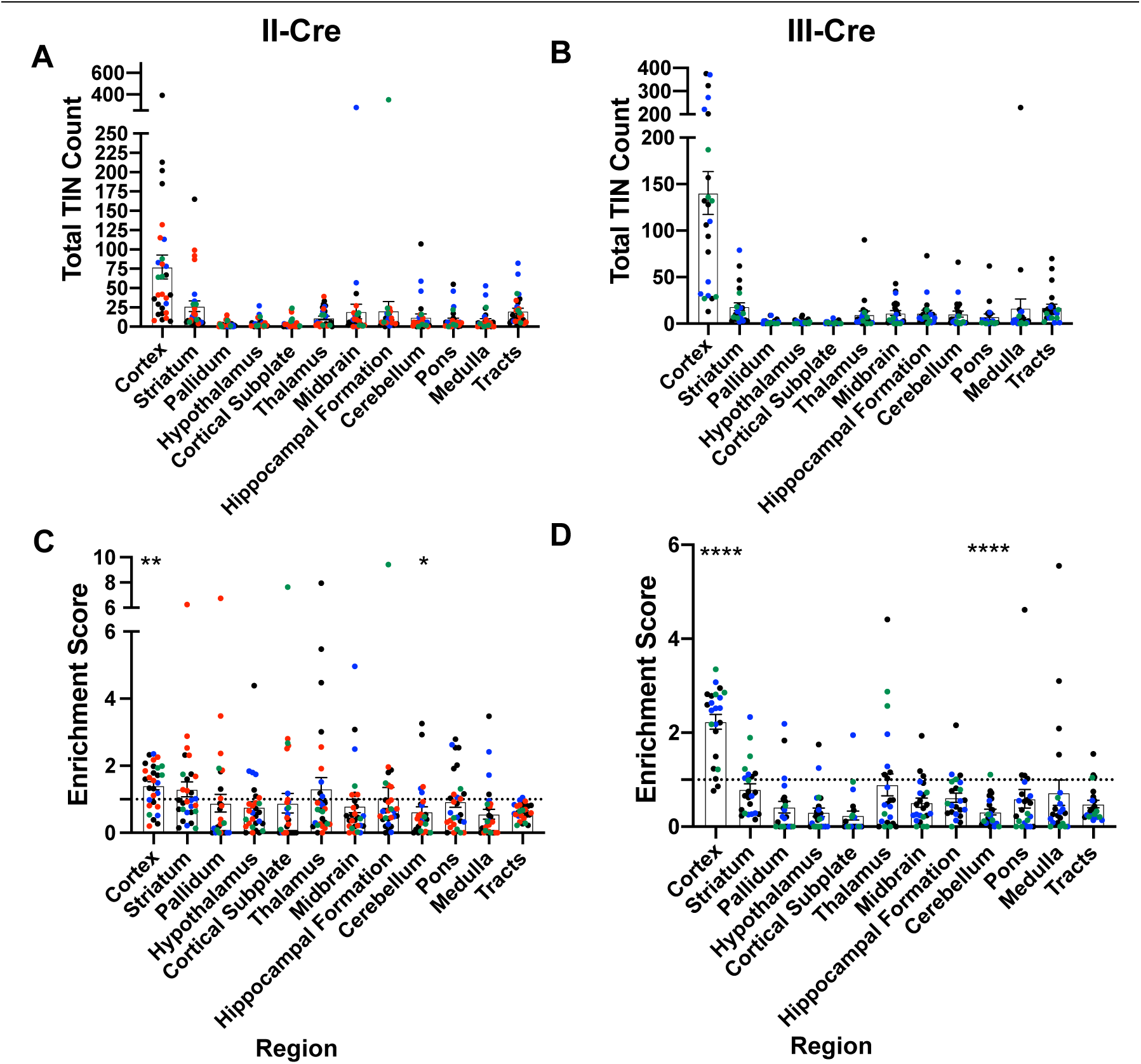
TINs show a predilection for the cortex at 3 weeks post infection. Cre-reporter mice were infected with II-Cre or III-Cre *Toxoplasma* parasites as indicated. Brains were harvested, sectioned, labeled, and quantified as previously described (Mendez, Potter et al., 2018). **(A, B)** Graphs of the absolute numbers of TINs mapped to 12 regions of the brain. **(C, D)** Graphs of TINs/region normalized to the size of the region. The dashed line is 1, the value at which TINs distribution would be considered proportional to the region size. Box ± bars, mean ± SEM. N = 16-20/sections/mouse. Individual colors denote animals from individual cohorts, N = 4-12 mice/cohort for II-Cre infected mice, 5-8 mice/cohort for III-Cre infected mice. * = p = 0.0170, ** = p = 0.0021 by one-sample t-test, **** = p ≤ 0.0001 P-values for all regions are in **Supplemental Table 2.**

As the cortex and striatum encompass a large proportion of the brain, the high number of TINs in these areas might simply be driven by their relatively size. To test this possibility, we produced an enrichment index using the following equation: 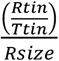, where *Rtin* is the TIN count for a given region (e.g. cortex), *Ttin* is the total TIN count from an individual mouse, and *Rsize* is the total area percentage of a specific region (e.g.% of brain that is cortex) (see **Supplemental Table 1** for area percentage for each brain region). As our method of sectioning does not allow us to collect the smaller sections of the cerebellum, we only included the cerebellar areas from sections 10-16 of the Allen atlas for normalization (consistent with the regions of the cerebellum which were analyzed). To align our sections with the Allen Institute sections, we also excluded the olfactory bulb (Mendez, Potter et al., 2018). Even with these exclusions, 85% of the original Allen atlas areas was used. If the TINs number in a given region is proportional to the size of the region, the normalization index will approximate 1, while regions enriched for TINs will have indices > 1, and regions devoid of TINs will have indices < 1. Using this normalization, we determined that— for both II-Cre and III-Cre infected mice— the cortex was the only region which was significantly enriched for TINs (II-Cre mice- index score 1.4 ± 0.1 p = 0.0021; III-Cre mice- index score 2.2 ± 0.2 p ≤ 0.0001) (**Figure 1 C, D**). For both groups of infected mice, within the cortex, the somatosensory, motor, and visual cortices had the highest enrichment score (**Supplemental Figure 2**). Finally, two regions— the cerebellum and the white matter tracts— showed a significant lack of enrichment in both II-Cre and III-Cre infected mice (**Figure 1 C, D**). **Supplemental Table 2** has the full list of enrichment scores and statistical analysis for the 12 listed regions.

Collectively these data show that, irrespective of infecting *Toxoplasma* strain, TINs are most commonly found in the cortex followed by the striatum, and relatively rarely found in the cerebellum. While TINs localization in the striatum is consistent with its size, the enrichment of TINs in the cortex and lack of TINs in the cerebellum are not accounted for by the relative size of these regions.

### TINs rarely co-localize with markers of inhibitory neurons

As the cortex and the striatum had the highest number of TINs, we decided to use these areas to determine if *Toxoplasma* targeted a specific neuron population. Neurons can be classified in many ways, including by neurotransmitter expression, physiology, and morphology. The simplest way is to classify neurons is as inhibitory or excitatory. Given that prior work suggested that *Toxoplasma* infection might specifically affect inhibitory neurons (Brooks et al., 2015), we first examined how often TINs co-localized with two common markers for inhibitory interneurons, calbindin and parvalbumin. We chose these makers rather than the pan-inhibitory marker glutamate decarboxylase (GAD) [GAD^+^ interneurons make up approximately 15-20% of all cortical neurons (Gentet et al., 2000; Lodato et al., 2015)] as GAD staining is altered in *Toxoplasma-*infected brain (Brooks et al., 2015; Carillo et al., 2020). Fortunately, parvalbumin labels 40-50% of GAD^+^ interneurons and calbindin labels approximately 30% of GAD^+^ interneurons (Hof et al., 1999; Keller et al, 2019). To quantify how often TINs were calbindin or parvalbumin interneurons, we labeled brain sections with antibodies against either calbindin or parvalbumin, and with antibodies against NeuN, a pan-neuronal marker. These labeled sections were then analyzed with confocal microscopy to identify TINs that co-stained with NeuN and either calbindin or parvalbumin. Within the cortex, irrespective of infecting strain, approximately 5% (4.8% ± 2.0 II-Cre, 3.9% ± 0.6% III-Cre) of TINs were calbindin^+^ (**Figure 2 A, B**), while none were parvalbumin^+^ (**Figure 2 C, D**). In the striatum, we observed that approximately 4% of II-Cre TINs were calbindin^+^ (4.4% ± 2.0%) and approximately 9% of III-Cre TINs were calbindin^+^ (9.4% ± 3.4%) (**Figure 2 E, F**). As in the cortex, no TINs co-localized with parvalbumin^+^ neurons in the striatum (N = 6-65 TINs/mouse, 4 mice for II-Cre; N = 30 and 84 TINs/mouse, 2 mice for III-Cre).

**Figure 2.**
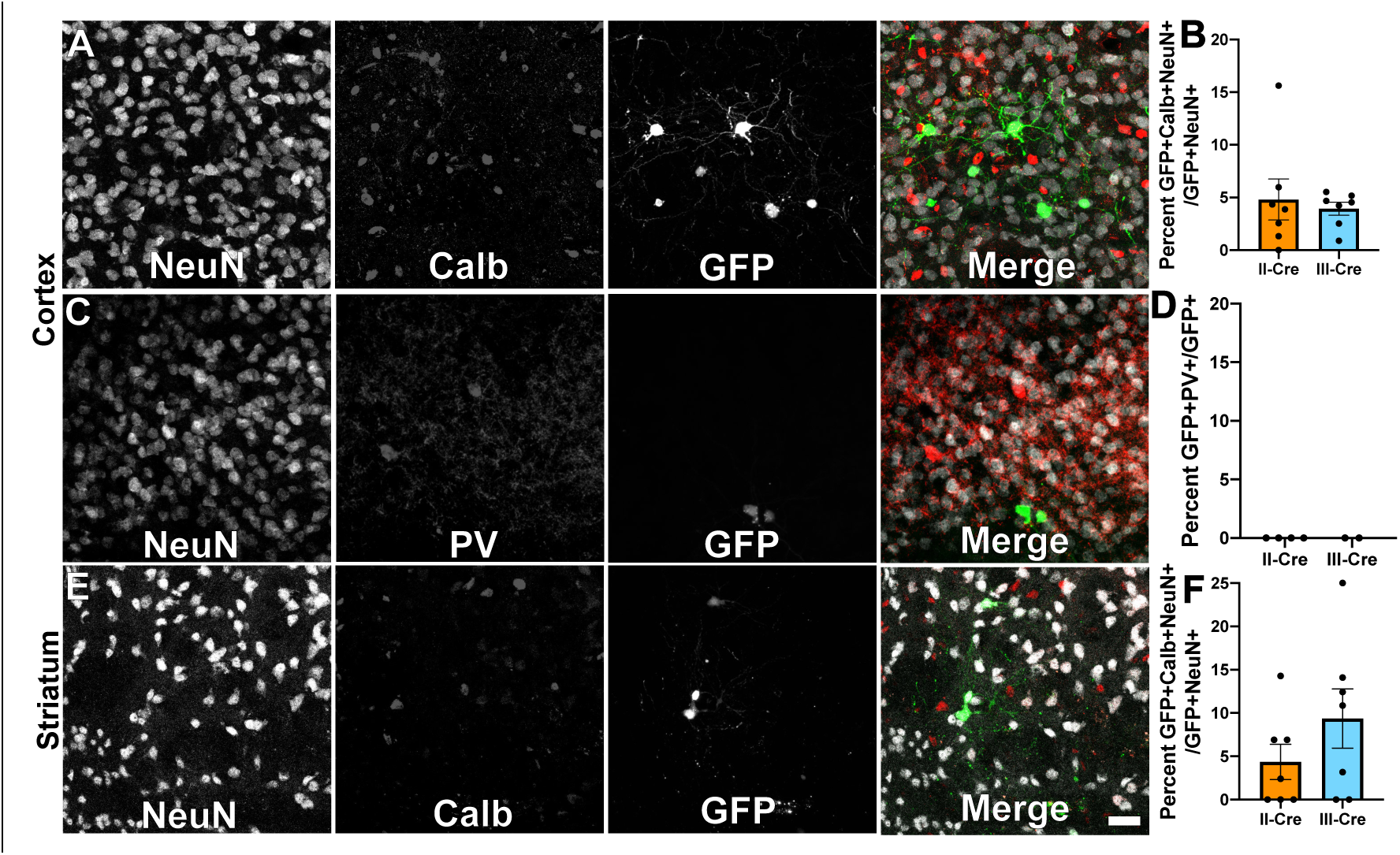
TINs rarely co-localize with inhibitory interneurons. Forty *μ*m brain sections from II-Cre or III-Cre infected mice were stained with either anti-calbindin or anti-parvalbumin antibodies. Stained sections were then imaged by confocal microscopy to determine co-localization between TINs (GFP^+^) and calbindin (Calb) or parvalbumin (PV) staining. **(A)** Representative images of a cortical region from a section stained with Calb. (**B**) Quantification of the percentage of TINs that co-localized with Calb staining. **(C)** As in (**A**) except the images are of a cortical region stained for PV. (**D**) As in (**B**) except for PV quantification. (**E**) as in (**A**), but in the striatum. For (**A, C, E**) Merge images, gray = NeuN, red = Calb or PV, and green = GFP. N = 9 sections/mouse, 7 mice/group for Calb, 2-4 mice/group for PV. (**B**) For II-Cre infected mice, 24-127 TINs/mouse were analyzed; for III-Cre infected mice, 254-503 TINs/mouse were analyzed. (**D**) For II-Cre 8-68 TINs/mouse and for III-Cre 281, 346 TINs/mouse, were analyzed. (**F**) For II-Cre infected mice, 3-290 TINs/mouse were analyzed; for III-Cre infected mice, 21-214 TINs/mouse were analyzed. No significant differences were identified between groups, Student’s T-test.

Collectively, these data suggest that *Toxoplasma* rarely interacts or injects cortical or striatal interneurons. Within interneurons, *Toxoplasma* appears to inject calbindin^+^ interneurons, not parvalbumin^+^ interneurons, an unexpected finding given the higher prevalence of parvalbumin^+^ interneurons.

### TINs colocalize with FoxP2, a marker of cortical excitatory neurons and medium spiny neurons

As only a small number of TINs co-localized with markers for inhibitory neurons, we next sought to determine if TINs co-localized with a marker for excitatory cortical neurons. For these studies, we selected the transcription factor FoxP2. While FoxP2 is primarily expressed by glutamatergic/excitatory neurons within layer 6 of the mouse cortex, some FoxP2^+^ excitatory neurons are also found in cortical layers IV and V (Arlotta et al., 2005; Lodato et al., 2015). In addition, FoxP2 expression occurs in medium spiny neurons in the striatum, allowing us to use a single marker for both cortical and striatal studies (Ferland et al., 2003; Lai et al., 2003). We quantified how often TINs were FoxP2^+^ by labeling and analyzing brain sections as above except now using antibodies directed against FoxP2. Within the cortex, irrespective of infecting *Toxoplasma* strain, approximately 58% (58.0% ± 3.8% II-Cre, 58.3% ± 2.3% III-Cre) of the analyzed TINs showed FoxP2 co-localization (**Figure 3 A, B**). In the striatum, we found that ~68% (68.1% ± 7.2%) of analyzed II-Cre TINs and ~ 83% (83% ± 4.1%) of analyzed III-Cre TINs showed FoxP2 co-localization (**Figure 3 C, D**).

**Figure 3.**
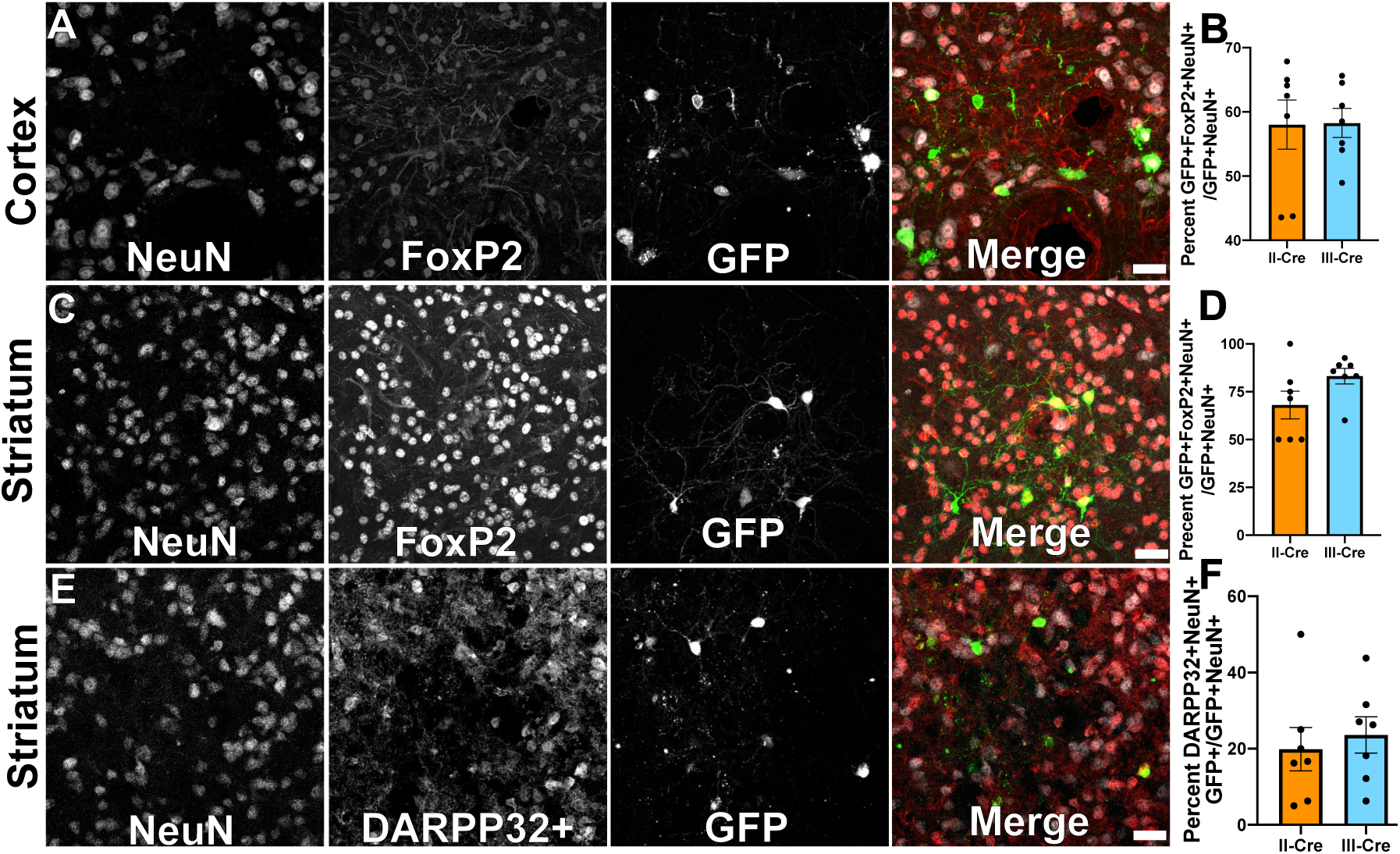
TINs co-localize with FoxP2 and DARPP32 staining. Brain sections from II-Cre or III-Cre infected mice were stained with anti-NeuN antibodies and anti-FoxP2 antibodies or anti-DARPP32 antibodies. DARPP32 staining was only done in the striatum. Stained sections were then analyzed by confocal microscopy to determine co-localization between TINs (GFP^+^) and FoxP2 or DARPP32 staining. **(A)** Representative image of a cortical region from a section stained as labeled. **(B)** Quantification of the percentage of TINs that co-localized with FoxP2 staining. (**C)** As in **(A)** except the images are of a striatal region. **(D)** As in **(B)** except for striatal TINs. **(E)** As in **(C)** except the tissue is stained with anti-DARPP32 antibodies (as labeled). (**F**) As in **(D)** except co-localization is between TINs and DARPP32 staining. **(A, C, E)** Merge images, gray = NeuN, red = FoxP2 or DARPP32, and green = GFP. Scale bar = 50*μ*m. **(B, D, F)** bars = mean ± SEM. For **(B,D, F)** N = 9 sections/mouse, 7 mice/group. (**B**) For II-Cre infected mice, a total of 24-127 TINs/mouse were analyzed; for III-Cre infected mice, 254-503 TINs/mouse were analyzed. (**D**) For II-Cre infected mice, a total of 28-68 TINs/mouse were analyzed; for III-Cre infected mice, 290-858 TINs/mouse were analyzed. **(F)** For II-Cre infected mice 6-237 TINs/mouse were analyzed; for III-Cre infected mice, 16-198 TINs/mouse were analyzed. No significant differences were identified between groups, Student’s T-test.

This high rate of co-localization between striatal TINs and FoxP2 suggested striatal TINs were medium spiny neurons (MSNs), which would be expected given that 90-95% of neurons in the striatum are MSNs (Ouimet et al., 1984; Ouimet & Greengard, 1990). As the identification of striatal TINs as MSNs had important implications for pursuing electrophysiology studies, we sought to further confirm that striatal TINs were MSNs by determined the rate of co-localization between striatal TINs and DARPP32^+^, another marker for MSNs (Ouimet et al., 1984; Ouimet & Greengard, 1990). We found that ~ 20% (19.9% ± 5.7%) or II-Cre striatal TINs and ~25% (23.6% ± 4.8) of III-Cre striatal TINs co-localized with DARPP32^+^ staining (**Figure 3E, F**).

Given the relatively low percentage of TINs co-localizing with DARPP32 and prior work suggesting *Toxoplasma* infection alters protein localization and expression (Cekanaviciute et al., 2014; Brooks et al., 2015), we sought to determine how striatal FoxP2 and DARPP32 staining differed between uninfected and infected mice, and if this staining differed by proximity to TINs. To determine the abundance of FoxP2 and DARPP32 neurons near TINs, in the images used for **Figure 3**, we quantified the number of neurons (NeuN^+^ cells) that co-localized with FoxP2 or DARPP32. To determine the overall abundance of FoxP2 and DARPP32 neurons, we again quantified the co-localization between neurons and FoxP2 or DARPP32, but this time used newly obtained, randomly distributed images from a subset of the mice analyzed for **Figure 3**. We found that the average number of FoxP2^+^ neurons in each image did not differ between images from either uninfected or infected mice, regardless of infecting *Toxoplasma* strain (**Supplemental Figure 3A**). Nor did the abundance vary in proximity to TINs (**Supplemental Figure 3B**). The average number of DARPP32^+^ neurons in each image also did not differ between random images from uninfected or infected mice (**Supplemental Figure 3C**). Conversely, in images taken in proximity to TINs, the number of DARPP32^+^ neurons showed a statistically significant decrease (**Supplemental Figure 3D**). In addition, we noted that, on average, more striatal neurons were FoxP2^+^ than DARPP32^+^, even in uninfected mice (**Supplemental Figure 3A, C**). To further test the possibility that FoxP2 identifies more striatal neurons than DARPP32, we stained tissues sections for both FoxP2 and DARPP32 and compared the co-localization between these proteins. Consistent with the findings in the single stains, only ~ 70% of FoxP2^+^ neurons stained for DARPP32 (69.6% ± 0.3% saline, 68.6% ± 0.6% II-Cre, 73.0 ± 4.0% III-Cre) (**Supplemental Figure 4A**), while 99% (99.7% ± 0.0% saline, 99.8% ± 0.0% II-Cre, 99.8% ± 0.1% III-Cre) of DARPP32^+^ neurons stained for FoxP2 (**Supplemental Figure 4B**).

Collectively these data suggest that striatal TINs are likely MSNs as they co-localize with markers of MSNs, especially FoxP2. The higher co-localization of TINs with FoxP2 versus DARPP32 is likely driven by the higher number of FoxP2^+^ neurons within the striatum as well as the sensitivity of DARPP32 expression/staining to disruption by infection or inflammation. Given that FoxP2 stains more striatal neurons than DARPP32, even in uninfected mice, FoxP2 may be a more sensitive marker for striatal MSNs than DARPP32.

### Passive electrophysiologic properties of bystander medium spiny neurons are relatively stable

The identification of striatal TINs as co-localizing with markers for medium spiny neurons (MSNs) offered us an unusual opportunity. MSNs make up 90-95% of dorsal striatal neurons (Mensah & Deadwyler, 1974; Graveland & DiFiglia M, 1985; Gerfen, 1992) and are highly characterized, including at the level of single cell electrophysiology *in situ* (i.e. patch-clamping in *ex vivo* slices). Targeting such a well-characterized, high frequency population would allow us to directly compare the electrophysiology of TINs and bystander neurons— striatal neurons in proximity to a TIN but not injected with *Toxoplasma* protein— as both groups of neurons would likely be MSNs. We reasoned that comparing TINs and bystander neuron physiology would allow us to determine the role of general inflammation in driving any changes we observed in TINs (i.e. if TINs and bystanders showed similar physiologic changes, then these changes are likely driven by the general inflammatory response to *Toxoplasma*, not from direct manipulation of TINs by *Toxoplasma* effector proteins.) For these studies, we chose to record from III-Cre infected mice because the type III strain is less virulent (meaning we could increase our inoculum as necessary), produces an overall increased frequency of TINs (**Figure 1A**), and shows higher rates of co-localization with the MSN marker FoxP2 (**Figure 3F**).

To ensure our capability to patch-clamp onto MSNs in *ex vivo* slices, we obtained thick brain slices from uninfected mice and performed patch-clamping using a standard protocol (Nisenbaum et al., 1994; Suter et al., 1999; Haubensak et al., 2010; Wang et al., 2019). As expected, in uninfected mice, dorsal striatum neurons showed classical MSN electrophysiologic properties such as a hyperpolarized resting membrane potential of ~ −80mV and a delayed first AP (**Figure 4A**). In addition, when we back filled a neuron showing the above characteristics and stained the section for DARPP32, we found the back-filled neuron co-localized with DARPP32 labeling (**Figure 4B-D**).

**Figure 4.**
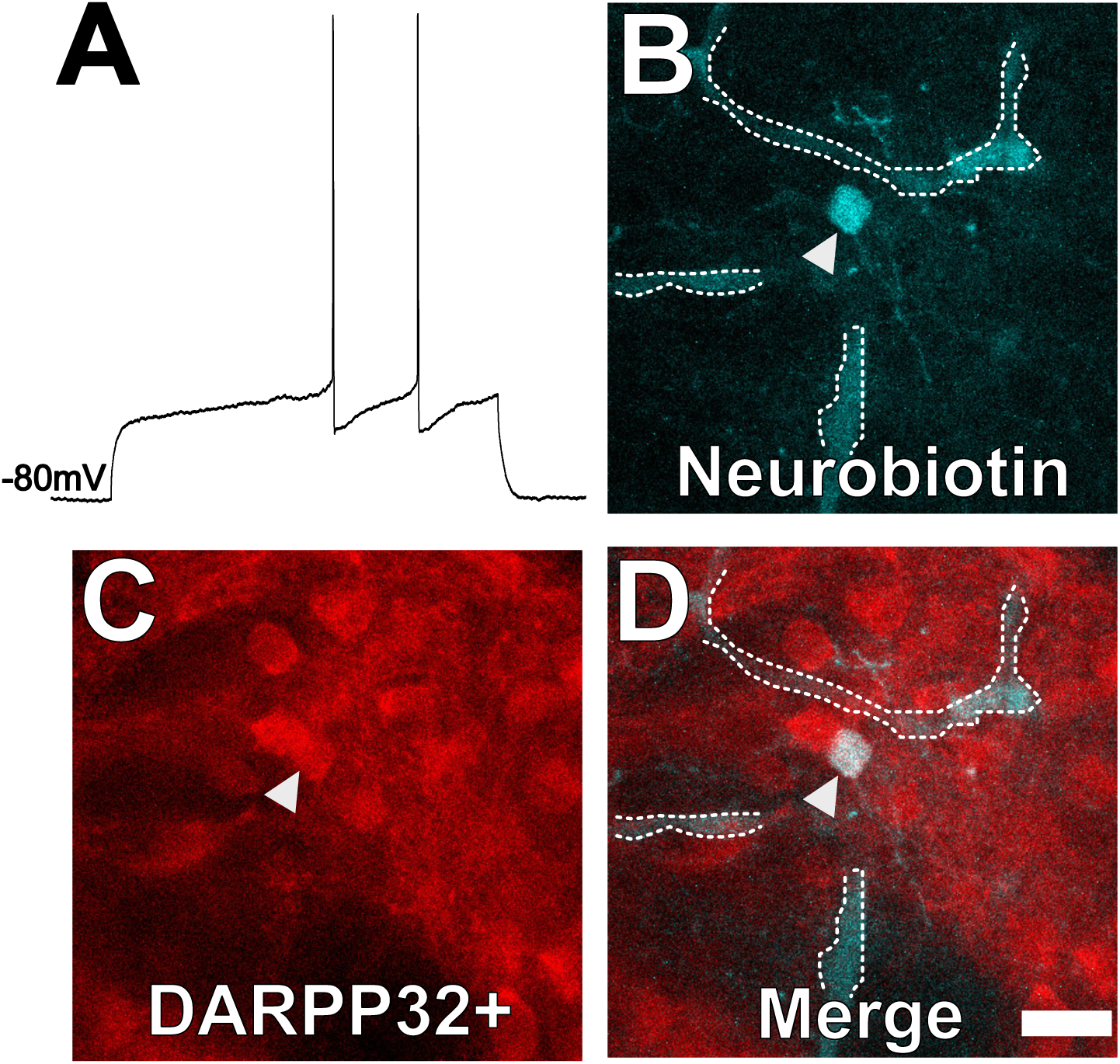
Neuron with electorphysiology of a MSN co-localizes with DARPP32 staining. (**A**) Sample tracing from the shown neuron. Note the hyperpolarized resting membrane potential of −80mV and the delayed time to the first action potential. (**B**) Image of patched neuron filled with neurobiotin. Arrowhead points to filled neuron. Dashed white line denotes biotin-filled blood vessels. (**C**) Image of DARPP32 staining. After recording and filling, the section was then fixed and counter labeled with anti-DARPP32 antibodies. (**D**) Merge of images **B** and **C**. **(B-D)** Section imaged on a Zeiss 880 confocal microscope. The shown images are a maximum projection of 12, 1*μ*m step images, from a 100*μ*m z-stack. Scale bar = 50*μ*m.

In infected mice, we electrophysiologically interrogated bystander MSNs and TINs, with bystander MSNs being approximately 30 – 100 *μ*m from a TIN. Bystander neurons had a resting membrane potential of −69.0mV ± 1.2, which is mildly depolarized compared to MSNs in uninfected mice (**Figure 5A**). Besides the elevated resting membrane potential of bystander neurons, the following electrophysiologic properties were equivalent when compared to MSNs from uninfected mice: action potential threshold, after hyperpolarization peak, action potential amplitude, delay to first action potential, and input resistance (**Figure 5B-G**).

**Figure 5.**
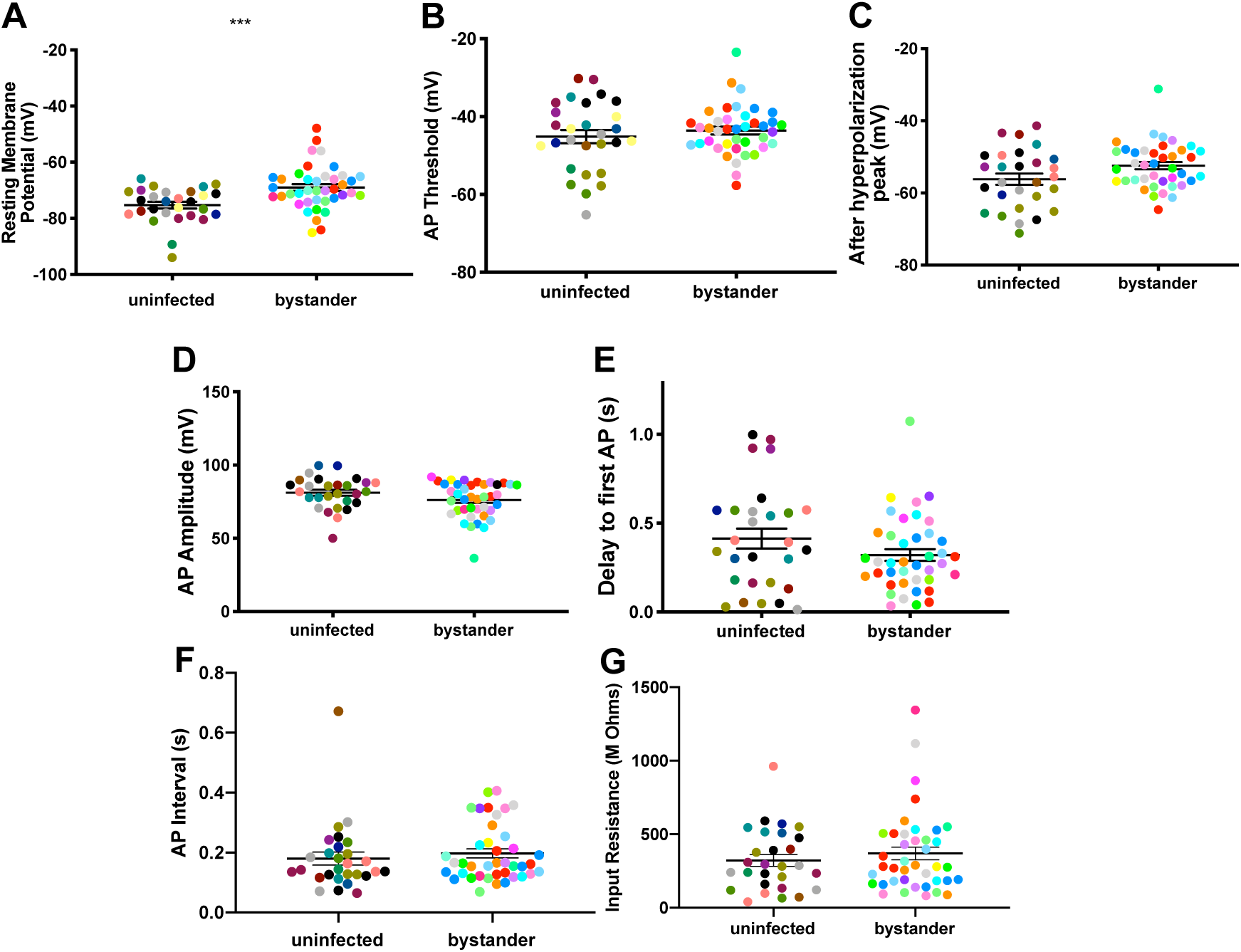
Bystander medium spiny neurons (MSNs) show similar electrophysiology as MSNs from uninfected mice. (**A**) Graph of the resting membrane potential of whole cell patch clamped MSNs in uninfected mice (−74.9mV ± 2.07) and bystander MSNs in infected mice (−68.7mV ± 4.0). (**B**) Action potential threshold, (**C**) after hyperpolarization peak, (**D**) action potential amplitude, (**E**) delay to first action potential, (**F**) action potential interval, and (**G**) input resistance in MSNs from uninfected mice and bystander MSNs in infected mice. Dots represent individual MSNs. Matching color dots denote cells from the same mouse. Uninfected MSNs, N = 28 cells recorded from 14 mice, 1-5 cells recorded/mouse. Bystander MSNs, N = 40 cells recorded from 18 mice, 1-5 cells recorded/mouse. Bars, mean ± SEM. p<0.001, Mann Whitney U-test.

Collectively these results indicate that though there is a mild change in the resting membrane potential of bystander neurons during infection, the infected/inflammatory microenvironment at this time point causes relatively little change to neurons that have not directly interacted with *Toxoplasma*.

### TINs show a highly depolarized membrane potential

We next evaluated TINs by following the same procedure as when we recorded from uninfected and bystander neurons, except that we used the expression of GFP to identify the TIN (**Supplemental Figure 5**). Unexpectedly, we were unable to properly patch onto most striatal TINs (i.e. we were unable to form a gigaseal). We were able to record from a total of 10 TINs, most of which had highly depolarized resting membrane potentials (~ −49.1 mV ± 4.5 mV) compared to either bystander or uninfected MSNs (**Figure 6A**). None of these TINs generated action potentials in response to the standard protocol, possibly because this highly depolarized membrane potential is too close to the threshold for firing an action potential (AP) (~45mV) in MSNs (**Figure 5B**). Several possibilities could most easily explain this highly depolarized membraned potential: i) the patched GFP^+^ cells were not neurons, ii) the patched cells were dead neurons, or iii) the patched cells were unhealthy neurons. To distinguish between these possibilities, when possible, we hyperpolarized the patched TINs, followed by attempts to generate action potentials. For 5 TINs, after we applied between −20 to −180pA of negative current, allowing us to achieve membrane potentials of ~ −55 to −66mV, we found that these TINs could generate APs (**Figure 6B**), definitively showing that they were neurons.

**Figure 6.**
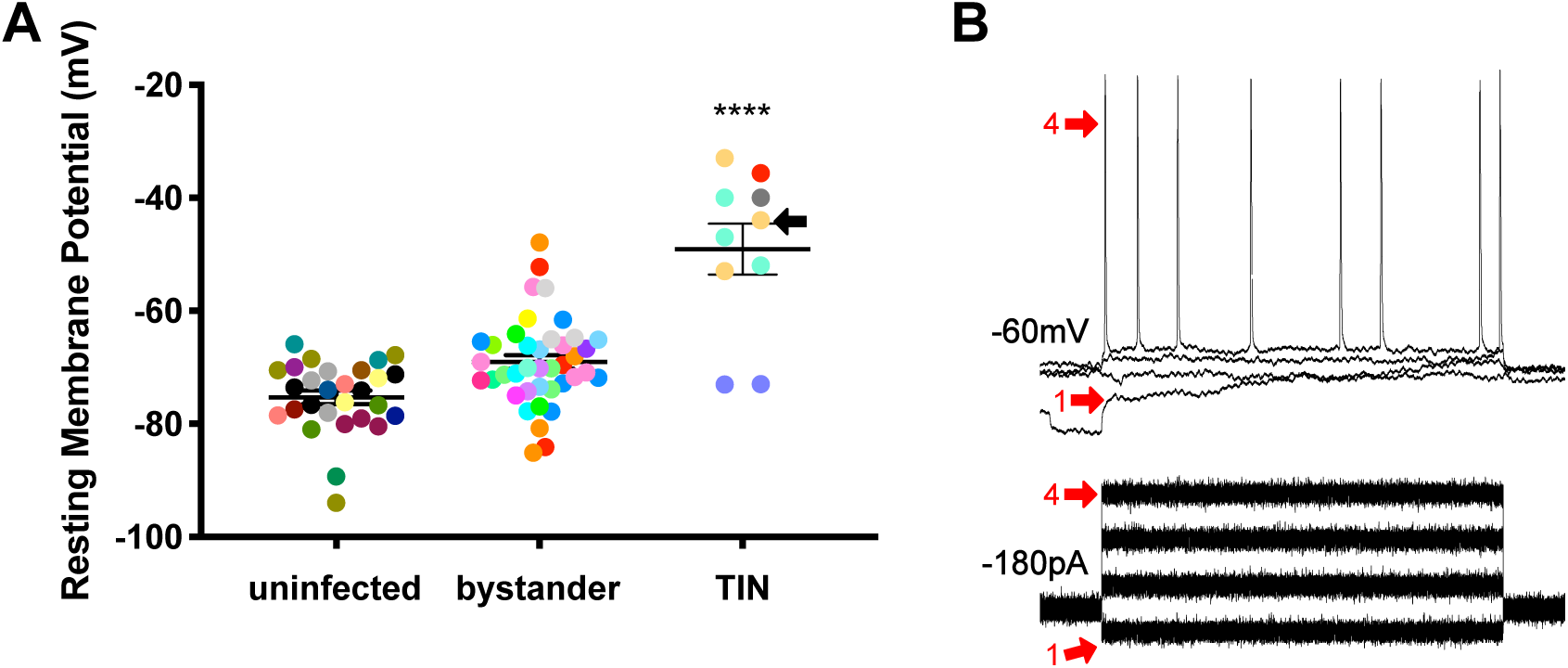
Unlike bystander neurons, TINs have highly abnormal electrophysiology. **(A)** TINs recorded resting potential mean = −49.1 ± 4.48 (SEM). **** = 0.001, one-way ANOVA with Tukey’s post-test. For TINs, N=10 cells recorded from 6 mice; uninfected and bystander described in **Figure 20**. Black arrow identifies TIN shown in **(B)**. **(B)***Top*: Example traces from TIN after hyper-polarization. *Bottom*: Visualization of a portion of the current injection protocol. The identified TIN was hyperpolarized to a resting membrane potential of −60mV by the injection of −180 pA of current. The red arrow denotes a single hyperpolarized step. The numbers (1, 4) match the voltage measurement (tracings) with the steps of increasing current. Note that an action potential is finally generated at the 4^th^ step of increasing current.

Given that such findings are consistent with the electrophysiology of sick or dying neurons (Lipton, 1999), we compared the total number of TINs in brain sections from mice infected with III-Cre parasites for 3 or 8 weeks. Consistent with the possibility that the electrophysiology reflects that TINs are sick/dying, brain sections from mice infected for 8 weeks showed an ~ 90% reduction in TINs compared to brain sections from mice infected for 3 weeks (**Figure 7**).

**Figure 7.**
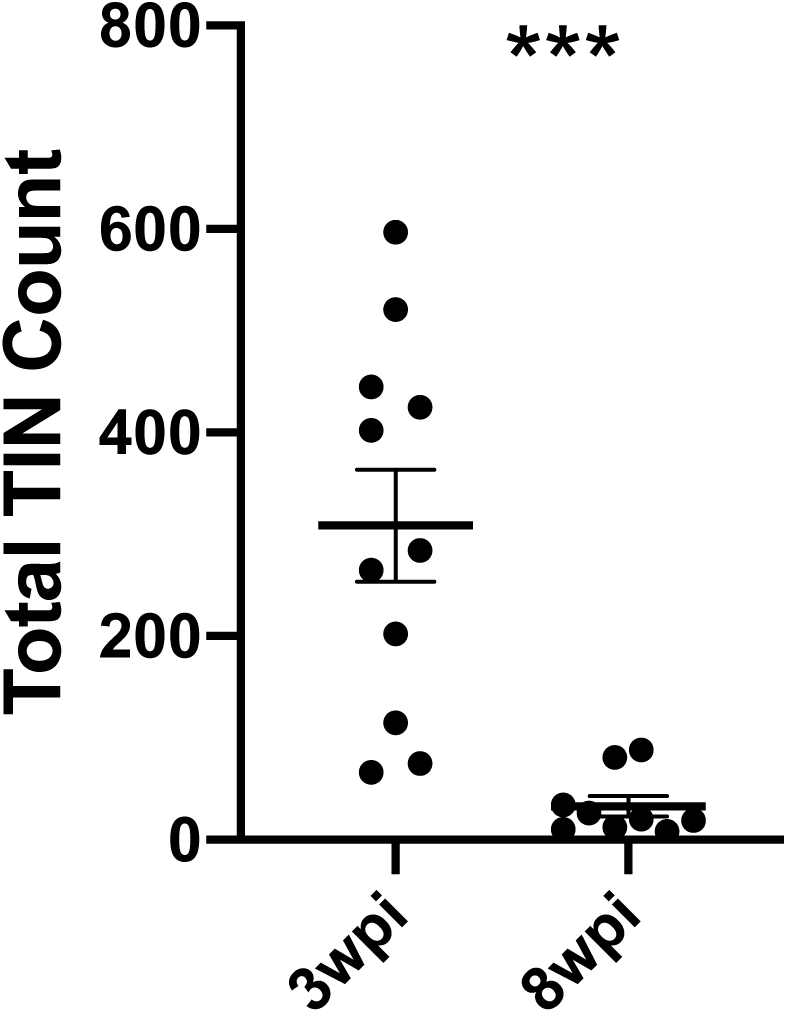
TINs numbers decrease by 8 weeks post infection. Mice were infected and, at 3 or 8 wpi, analyzed as in **Figure 12** except that total TINs numbers were quantified. Bars, mean ± SEM. ***p ≤ 0.0001, Mann-Whitney U test. 16-20 sections/mouse, N=9-11 mice/time point.

Collectively, our data suggest that neurons that directly interact with parasites are highly altered at the level of electrophysiology and most will die over a relatively short period. On the other hand, bystander neurons, which are in a similar inflammatory environment, show very little change in terms of electrophysiology. Together these data suggest that neuron pathology is extremely targeted, even in a highly inflammatory setting.

## Discussion

In this study, we leveraged the *Toxoplasma*-Cre system to extend our understanding of neuron-*Toxoplasma* interactions. Across multiple cohorts of mice and 2 *Toxoplasma* strains, our data show that *Toxoplasma*-injected neurons (TINs) consistently localize to the cortex and striatum. For the cortex, TINs numbers are higher than would be expected even after accounting for the size of the cortex. Within the cortex and striatum, we found little evidence of *Toxoplasma* targeting a specific neuron subtype, as TINs co-localize with markers for highly abundant neurons in a region (e.g. medium spiny neurons (MSNs) in the striatum). Interestingly, when we used whole cell patch clamping in *ex vivo* slices to assess both TINs and bystander neurons— neurons in the same infected milieu but not injected with parasite protein— we found discrepant electrophysiology. Bystander neurons showed mild changes in resting membrane potential and no changes in other passive firing properties, while, TINs had highly depolarized resting membrane potentials and did not fire action potentials unless artificially hyper-polarized. These data suggest that TINs, but not bystander neurons, are unhealthy and potentially dying, a possibility confirmed by a drastic decrease in TINs numbers at a later time point of infection. Collectively, these data offer an unprecedented look into the neuron-*Toxoplasma* interface, raising new questions about why the cortex is particularly vulnerable to *Toxoplasma* infection and the evolutionary advantage of the death of neurons injected with *Toxoplasma* protein.

One of the major advances of this work is how we mapped neuron-*Toxoplasma* interactions. Most prior studies have utilized a single *Toxoplasma* strain; relatively few animals (or more animals but relatively few sections/animal); and were limited to measuring cyst location. On the other hand, we mapped TINs, which outnumber cysts by over 20-fold (Koshy et al., 2012) and show uniform distribution of GFP, allowing us to map neuron cell bodies or somas which is the traditional method for identifying neuron anatomic location. Furthermore, our semi-automated method significantly reduces the time and effort required to perform this mapping (Mendez, Potter et al., 2018), making it feasible to map TINs in mice infected with 2 genetically distinct *Toxoplasma* strains (type II and type III) across multiple cohorts of mice and using 16-19 sections/mouse. These robust methods offer confidence that our findings are not unduly influenced by a single outlier mouse; a cohort effect; or insufficient sampling (i.e. using low numbers of tissue sections or animals in a process that is highly heterogenous). Our findings are consistent with several aspects of the two most extensive cyst mapping studies (Berenreiterova et al., 2011; Boillat, et al., 2020): all three studies found high levels of inter-mouse variability; ii) identified the cortex— and especially the motor, somatosensory, visual cortical regions— as showing enrichment for *Toxoplasma* presence (TINs or cysts) even when accounting for region size; and iii) determined that the cerebellum shows relatively little *Toxoplasma* presence, especially when accounting for size. These consistencies suggest that regional cyst burden is driven by differences in the number of neuron-parasite interactions, not by differences in regional innate immune responses as has been suggested for West Nile Virus (WNV) infection (Cho et al., 2013). In addition, these regional differences are consistent with the largest studies in AIDS patients with toxoplasmic encephalitis, in which more lesions are identified in the cortex and the basal ganglia compared to the cerebellum (Arendt et al., 1999; Porter & Sande, 1992), highlighting the relevance of the mouse model to human disease. Currently, we do not know what drives these regional differences in tropism, though one possibility is differences in the transmissibility of *Toxoplasma* across the vasculature that supplies the cortex and striatum/basal ganglia versus the cerebellum (anterior versus posterior circulation respectively), a possibility also suggested for WNV (Daniels et al., 2017).

A second advance in our study is the ability to identify the type of neuron injected by *Toxoplasma*. The finding that TINs are comprised of common neuron subtypes in the deep cortex and striatum (e.g. FoxP2^+^ neurons) suggests that *Toxoplasma* injects (and presumptively infects) in proportion to the neuron subtypes in a given region. One exception to this rule is the lack of TINs that also express parvalbumin. Parvalbumin-expressing neurons make up approximately 40-50% of the cortical GABAergic (inhibitory) neurons and are found in all deeper layers of the cortex, while calbindin-expressing neurons are less frequent, especially in layers IV-VI of the cortex (Hof et al., 1999; Tremblay et al., 2016). One possibility for the lack of parvalbumin^+^ TINs is a decrease or change in parvalbumin expression/location as seen for DARPP-32 (**Supplemental Figure 3**) or GAD-67 (Brooks et al., 2015). Another more exciting possibility is that parvalbumin^+^ neurons have fewer interactions with *Toxoplasma* because they have a high concentration of perineural nets— extracellular matrix that surrounds different parts of the neuron (Baker et al., 2017)— blocking *Toxoplasma*’s ability to contact parvalbumin^+^ neurons. Future studies will define the factors that determine which neurons interact with parasites.

A final advance in this work is the electrophysiology comparing TINs and bystander neurons. To the best of our knowledge, these studies are the first in any neurotropic infection to compare the *in situ* electrophysiology between microbe-interacted (or infected) neurons versus those neurons in close proximity but without direct microbial interactions. The utility of comparing TINs and bystander neurons is to address what physiologic effects are driven by the general neuroinflammatory response versus those changes driven by neuron-parasite interactions. Given the relatively common understanding that microglia will strip synapses off infected and uninfected neurons in the inflamed brain (Vasek et al., 2016; Di Liberto et al., 2018), we found it remarkable that bystander MSN physiology showed only minor changes compared to MSNs from uninfected mice while TINs showed highly abnormal electrophysiology (**Figure 6**). On possible explanation for the minimal changes seen in bystander neurons is that neurons that have not directly interacted with *Toxoplasma* undergo compensatory mechanisms that allows these neurons to avoid drastically changing, since a dramatic response would be detrimental to the entire local circuitry. Another possibility is that the effect of the neuroinflammatory response is confined to a small area near TINs, leaving these more distant bystander neurons relatively spared. Either explanation would preserve circuit function at a global level and potentially allow for relatively quick reversal of these physiologic changes once the inflammatory response subsides. Such possibilities are consistent with the relative lack of behavioral abnormalities seen in *Toxoplasma*-infected mice who have cleared both the infection and the immune response (McGovern et al., 2020).

TINs, on the other hand, show highly abnormal electrophysiology, only firing action potentials after artificial hyperpolarization. This remarkable discrepancy between TINs and bystander physiology suggests that whatever is causing the abnormal TIN’s physiology is specific to neurons injected with *Toxoplasma* protein. Such TINs-restricted changes could arise from direct manipulation of TINs by the injected parasite proteins or selective microglial/macrophage recognition and phagocytosis of TINs, consistent with what was observed in lymphocytic choriomeningitis virus infection (Di Liberto et al., 2018). Another possibility, as noted above, is that the immune response is tightly regulated and confined, such that the destruction leveled by this response is limited to a very small area around TINs. Such a possibility is consistent with prior work showing that partial abrogation of astrocytic TGF-β signaling led to de-regulation of the CNS immune response to *Toxoplasma* and higher levels of neuronal loss without changing the CNS parasite burden (Cekanaviciute et al., 2014). Finally, the ~ 90% decrease in TINs by 8 wpi questions dogma that the long-lived, rarely regenerated nature of neurons demands that neuron preservation trump immune cell cytolysis of infected neurons (Binder & Griffen, 2001; Patterson et al., 2002; Miller et al., 2016). This finding of TINs death is particularly intriguing because prior work suggests that >90% of these TINs are expected to be uninfected (Koshy et al., 2012). One possibility is that these *Toxoplasma*-injected neurons are so abnormal (from either direct manipulation by parasites or because of neuron-immune cell interactions) that global brain function is better preserved with the removal of these neurons. Future studies will determine what factors lead to TINs abnormal physiology and how and why TINs are cleared from the CNS.

## Materials and Methods

### Ethics statement

All mouse studies and breeding were carried out in accordance with the Public Health Service Policy on Human Care and Use of Laboratory Animals. The protocol was approved by the University of Arizona Institutional Animal Care and Use Committee.

### Parasite maintenance

Parasites were maintained via serial passage through human foreskin fibroblasts using DMEM supplemented with 2 mM glutagro, 10% fetal bovine serum, and 100 I.U./ml penicillin/ 100 *μ*g/ml streptomycin. The type II (Prugniaud) and type III (CEP) strains used have been engineered to express mCherry and Cre recombinase and have been previously described (Koshy et al., 2010; Koshy et al., 2012). For simplicity, these strains are denoted as II-Cre and III-Cre.

### Mice

Mice used in these studies are Cre-reporter mice that express a green fluorescent protein (GFP) only after the cells have undergone Cre-mediated recombination (Madisen et al., 2010). Mice were purchased from Jackson Laboratories (stock # 007906) and bred in the University of Arizona Animal Center.

### Infections

Mice were infected at 2-3 months of age via intraperitoneal (IP) injection with freshly syringe-released parasites (**Figure 1, 2, 3**). Mice were inoculated with 10,000 parasites in 200*μ*l of USP grade PBS for both II-Cre and III-Cre strains. To increase the reliability of electrophysiology studies, mice were infected at 5 to 6 weeks of age with III-Cre parasites only.

### Tissue preparation

For the localization studies, at 3 weeks post infection (wpi) animals were sedated with a ketamine/xylazine cocktail, intracardially perfused with saline followed by 4% paraformaldehyde, after which brains were harvested. Sections were then prepared as previously described (Mendez, Potter et al., 2018). In brief, left and right brain hemispheres were isolated, drop-fixed in 4% PFA followed by cryopreservation in 20% sucrose. Forty-micron thick sagittal sections were generated using a freezing sliding microtome (Microm HM 430). Sections were then stored as free-floating sections in cryoprotective media (0.05 M sodium phosphate buffer containing 30% glycerol and 30% ethylene glycol) until labeling procedure was to be done. Sections for localization studies were sampled every 200*μ*m (**Figures 1, Supplemental Figure 2**), while for co-localization studies (**Figures 2,3**), sections were sampled every 400 *μ*m.

### Immunohistochemistry

To ensure adhesion of tissue onto slides for localization studies, tissue was allowed to air-dry onto slides overnight then heated on a slide warmer for 40 minutes at 34°C. Then tissue was dehydrated using increasing then decreasing concentrations of 50%, 75%, 95%, and 100% ethanol. Slides were washed with TBS, peroxidase inactivated (3%H_2_O_2_/10% methanol), permeabilized (0.6% Triton X-100), blocked (1.5% BSA and 1.5% goat serum), and incubated with Rabbit anti-ZsGreen (Clontech, Cat. No. 632474, 1:10,000) for 15-18hrs at room temperature. Next, slides were incubated in Goat anti-rabbit polyclonal biotinylated conjugated antibody (Vector Labs, Cat. No. BA-1000, 1:500) for 2 hrs, incubated with avidin-biotin complex kit (ThermoFischer Scientific, 32020) for 2 hrs and visualized with a 3,3’-diaminobenzidine kit (Vectastain, Vector Labs, SK-4100). Sections were then counterstained with cresyl violet for Nissl labeling.

For co-localization studies sections were labeled free-floating, 8-10 sections per mouse. Sections were washed with PBS, permeabilized with 0.3% Triton-X, then permeabilized and blocked with 0.3% Triton-X and 5% goat serum (Jackson Immuno). For primary antibodies, the following were incubated for 15-18hrs at 4°C in PBS with 0.3% Triton-X and 5% goat serum: mouse biotin conjugated anti-NeuN B-clone A60 (MAB377, Millipore, 1:500), rabbit anti-calbindin (C2724, Sigma-Aldrich, 1:500), rabbit anti-parvalbumin (ab11427, abcam, 1:1000), rabbit anti-FoxP2 (ab16046, abcam, 1:1000), rabbit anti-DARPP32 (ab40801, abcam 1:500). Following incubation in the appropriate antibody, sections were then incubated for 2 hours in the following secondaries: goat anti-rabbit IgG Alexa Fluor 568 (A11011, Invitrogen, 1:500), Cy5 streptavidin (SA1011, Invitrogen, 1:500), and all sections were counterstained with 4′,6-diamidino-2-phenylindole (DAPI, ThermoFisher, 1:1000).

For filled medium spiny neurons (MSNs), sections were processed as discussed below. For staining, sections with filled MSNs were permeabilized with 0.6% Triton-X in PBS, then incubated with DARPP32 antibody in 5% goat serum, 0.6% Triton-X in PBS for 48hrs at 4°C. Sections were washed then incubated with Cy5 streptavidin (1:500) and goat anti-rabbit Alexa Fluor 568 (1:500) for 24hrs at 4°C.

### Electrophysiology of medium spiny neurons

Mouse brain slice electrophysiology recording was performed as described (Suter et al., 1999; Nisenbaum et al., 1994; Dorris et al., 2014; Willett et al., 2016; Haubensak et al., 2010). Mice were sacrificed with CO_2_, after which the descending aorta was clamped and the mice were intracardially perfused with ice cold oxygenated artificial cerebral spinal fluid (ACSF) containing: 126 mM NaCl, 1.6 mM KCl, 1.2 mM NaH_2_PO_4_, 1.2 mM MgCl_2_, 2.4 mM CaCl_2_, 18 mM NaHCO_3_, 11 mM glucose. After the olfactory bulbs and the cerebellum were removed, the brain was transferred to a vibratome stage that contained ice-cold ACSF oxygenated with carbogen (95% O_2_ balanced with CO_2_). 200 *μ*m coronal sections were then generated with the vibratome (Leica, VT1000S). Brain slices were immediately transferred to oxygenated NMDG-HEPES recovery solution (93 mM NMDG, 2.5 mM KCl, 1.2 mM NaH_2_PO_4_, 30 mM NaHCO_3_, 20 mM HEPES, 25 mM Glucose, 5 mM sodium ascorbate, 2 mM thiourea, 3 mM sodium pyruvate, 10 mM MgSO_4_, 0.5 mM CaCl_2_, 300–310 mOsm, titrated with 10 N HCl to adjust pH to 7.3–7.4) and were allowed to recover for 15 min at 32–34°C. After the recovery portion, brain slices were then transferred to room temperature oxygenated ACSF for 1hr. After the hour rest period, recordings were performed in a rig equipped with a MultiClamp 700B, a Digidata 1550A1 (Molecular Devices), and a fluorescent microscope (Olympus BX51) that was used to visualize sections and identify TINs. Patch pipettes were pulled with P-97 Sutter micropipette puller to achieve a resistance of 8–16 MΩ, after which they were filled with an intracellular solution (135 mM potassium gluconate, 5 mM EGTA, 0.5 mM CaCl_2_, 2 mM MgCl_2_, 10 mM HEPES, 2 mM MgATP, and 0.1 mM GTP, pH 7.3–7.4, 290–300 mOsm). Recording data were sampled at 10 kHz, filtered at 3 kHz, and analyzed with pCLAMP10.7. For bystander neurons and neurons in uninfected mice, the classification as a medium spiny neuron was based on the firing pattern in response to the current injections as described previously (Schmidt & Perkel, 1998; Farries & Perkel, 2000; Luo et al., 2001; Ding & Perkel, 2002; Ding et al., 2003, Willet et al., 2016). For filling experiments, once data from a 200 *μ*m section were recorded, the recorded neurons were filled with 0.6% neurobiotin (SP1120, Vector Labs). Filled sections were then stored in 4% paraformaldehyde for 24 hrs followed by cryopreservation in cryoprotective media until stained as described above.

### Microscopy and quantification

Slides for localization data were imaged on a Leica DMI6000 with a motorized stage, using Leica Application Suite X (LAS X) at 10x magnification. Base background subtraction and white balance was maintained throughout individual cohorts. Image stitching was done automatically through LAS X with a 10% overlap and images stored as Leica Image File Format (lif). Lif images were then processed on a custom MATLAB code for image transformation onto the Allen Institute reference atlas, followed by semiautomated quantification, as described in Mendez, Potter et al., 2018. Animals were excluded due to having the same or fewer TINs compared to saline injected animals (~15 TINs) and via ROUT outlier tests. **Supplemental Figure 1** has all mice included and includes 100% of the Allen atlas for the enrichment index. For co-localization studies, 8-10 sections/mouse were analyzed. Images were captured on a Zeiss LSM880 inverted confocal microscope (Marley Microscopy Core, UA). Images were captured with a 20x objective, with 1x PMT zoom, all settings were the same across specific staining runs. Autofocus was used to capture a z-stack that would capture from edge to edge of the section.

### Statistics

In **Figure 1**, data were analyzed with a one-sample t-test to compare the enrichment score to a value of 1, which would indicate a random distribution of TINs in a given region of the brain. For **Figure 2–3**, the data were analyzed with a two-tailed t-test to test for differences between II-Cre and III-Cre infected groups. For **Figure 5, 7** the data were analyzed with a Mann Whitney two-tailed t-test. All statistics were done via Prism statistical software (v8.4.2, Graphpad).

## Acknowledgments

The authors would like to thank the laboratory of Dr. Haijiang Cai for initial electrophysiology training and equipment usage. We would also like to thank Dr. David Perkel for helpful discussions about the electrophysiology data. Additional thanks to Patricia Jansma and Doug Cromey for imaging training. Finally, thank you to the entire Koshy lab for helpful discussions.

## Funding

AAK:
Arizona Biomedical Research Center (ADHS14-082991)
NINDS R01 NS095994
The BIO5 Institute, University of Arizona

OAM:
NINDS F99 NS108514

EFM:
NINDS R25 NS076437 High School Student NeuroResearch Program (HSNRP)

The funders had no role in study design, data collection and interpretation, or the decision to submit the work publication.

---

**Supplemental Figure 1.**
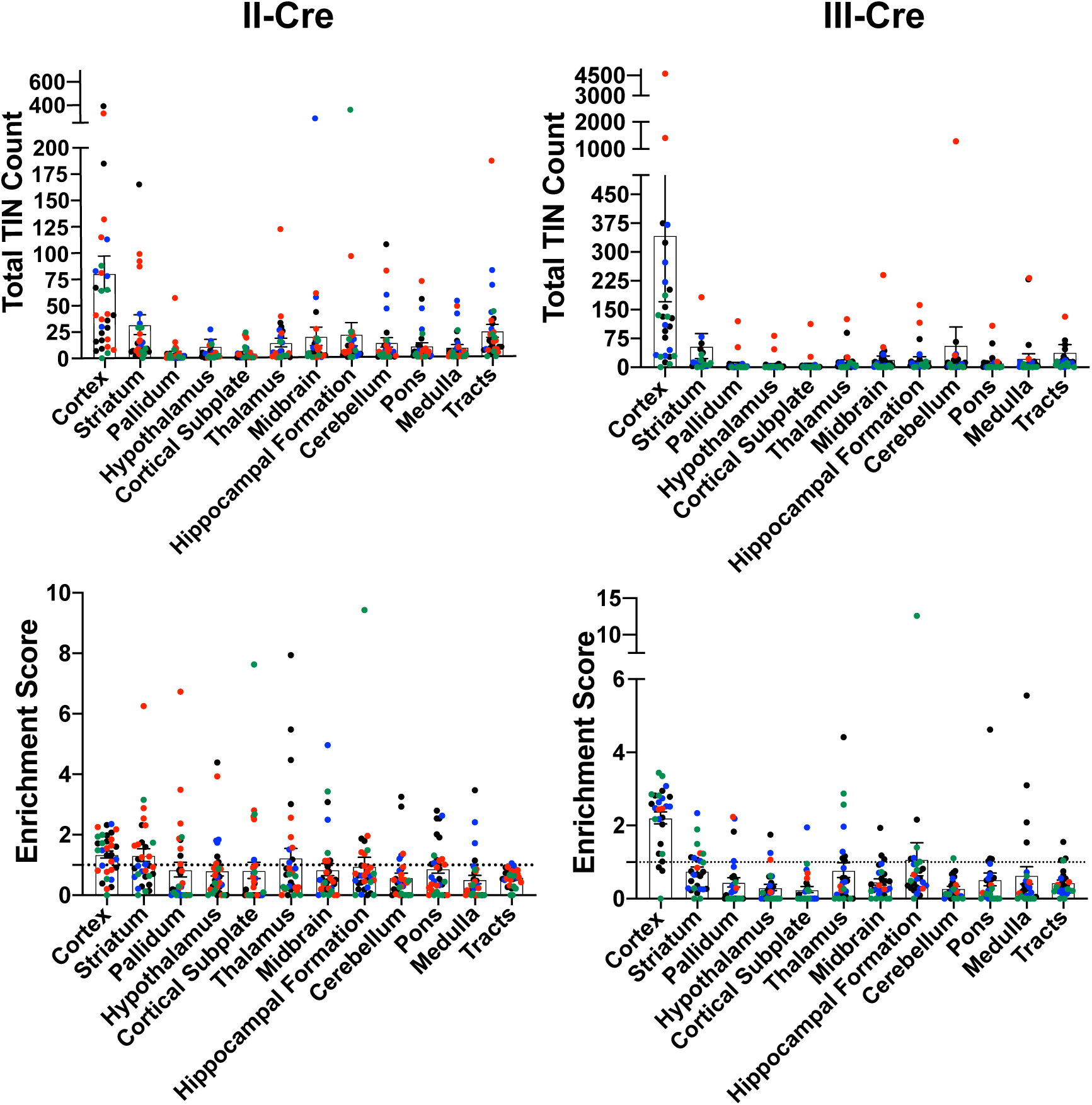
Original data set before exclusions. Cre-reporter mice were infected with II-Cre or III-Cre *Toxoplasma* parasites as indicated. Brains were harvested, sectioned, labeled, and analyzed as described in **Figure 1**. **(A, B)** Graphs of the absolute numbers of TINs mapped to 12 regions of the brain. **(C, D)** Graphs of TIN numbers/region normalized to the size of the region. The dashed line indicates the value at which TINs distribution would be considered random and appropriate for the region size. Data presented is before the exclusion of any mice. Exclusion was determined by i) the total number of TINs being less than or equal to the total GFP^+^ CNS cells in saline injected mice or ii) identification as an outlier via the ROUT method.

**Supplemental Figure 2.**
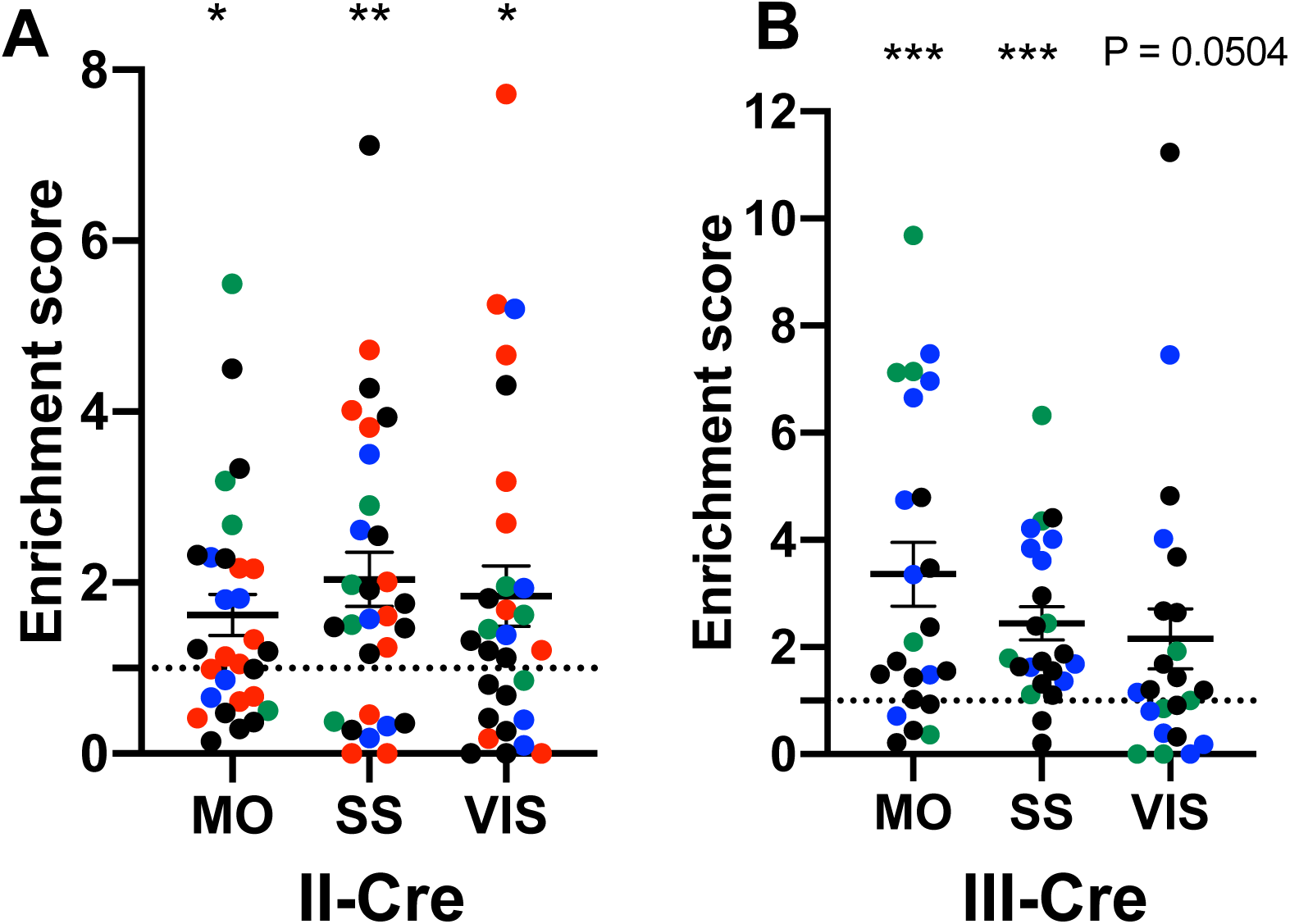
The visual, somatosensory, and motor cortices are highly enriched cortical regions containing TINs. **(A)** Graph showing the normalized distribution of cortical TINs from II-Cre infected mice in motor (MO), somatosensory (SS), and visual (VIS) cortices. **(B)** As in **(A)** except for III-Cre infected mice. * p ≤ 0.05, ** p=0.01, *** p ≤ 0.0005, by one-sample t-test. Individual colors denote mice from the same cohort.

**Supplemental Figure 3.**
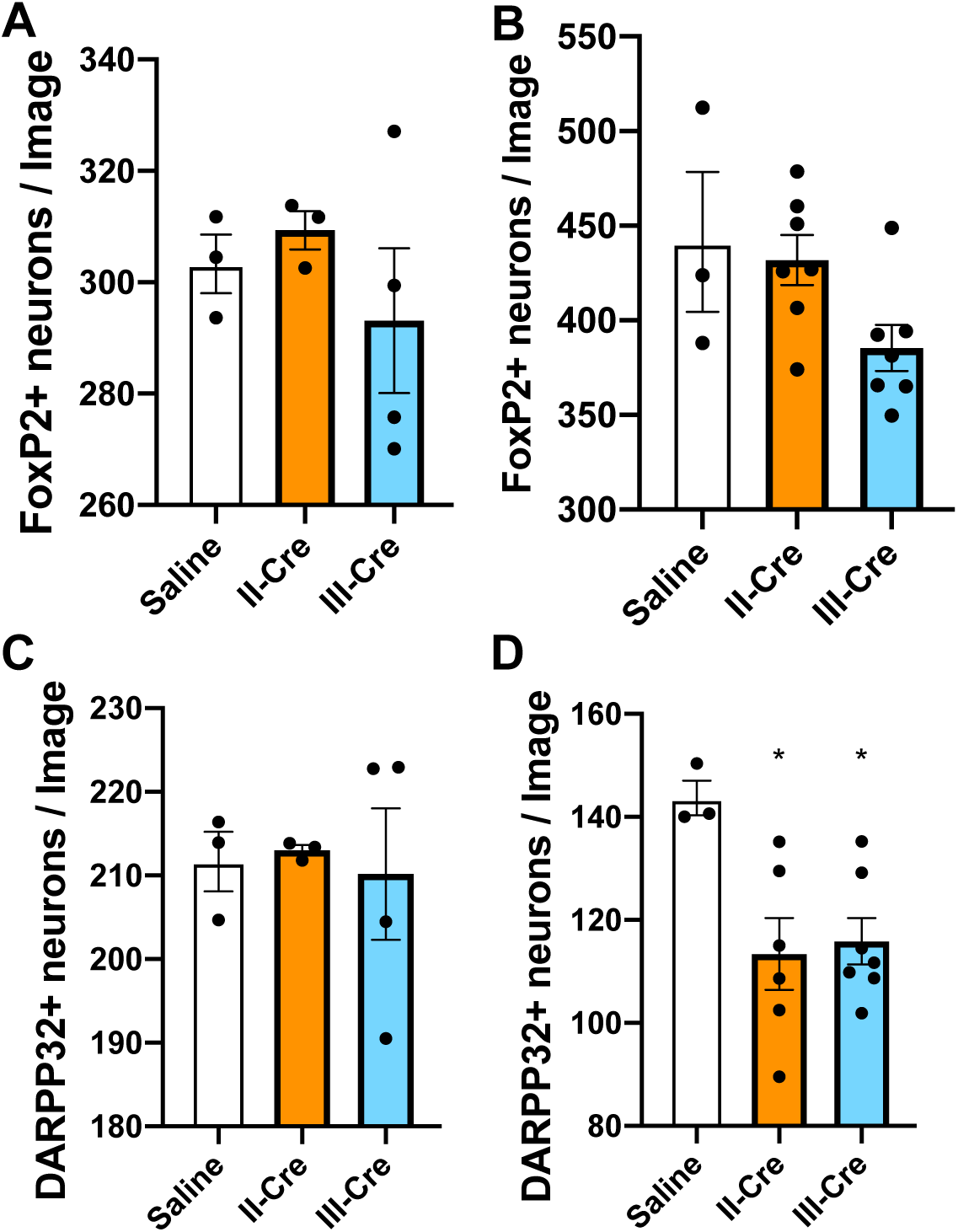
Quantification of FoxP2^+^ or DARPP32^+^ neurons in the striatum. Brain sections from 3 wpi saline or II-Cre or III-Cre inoculated mice were stained with anti-NeuN antibodies and anti-FoxP2 or anti-DARPP32 antibodies. Stained sections were then analyzed for the number of FoxP2+ or DARPP32+ neurons. **(A)** Average number of FoxP2^+^ neurons/image when randomly sampling the striatum **(B)** Average number of FoxP2^+^ neurons when sampling near TINs. **(C)** As in **(A)**, but for DARPP32^+^ neurons (random sampling). **(D)** As in **(B)** but for DARPP32^+^ neurons (near TINs). bars = mean ± SEM. N = 9 sections/mouse, 31-44 images/mouse, 3-7 mice/group. * p < 0.05, one-way ANOVA with Dunnett’s post-test.

**Supplemental Figure 4.**
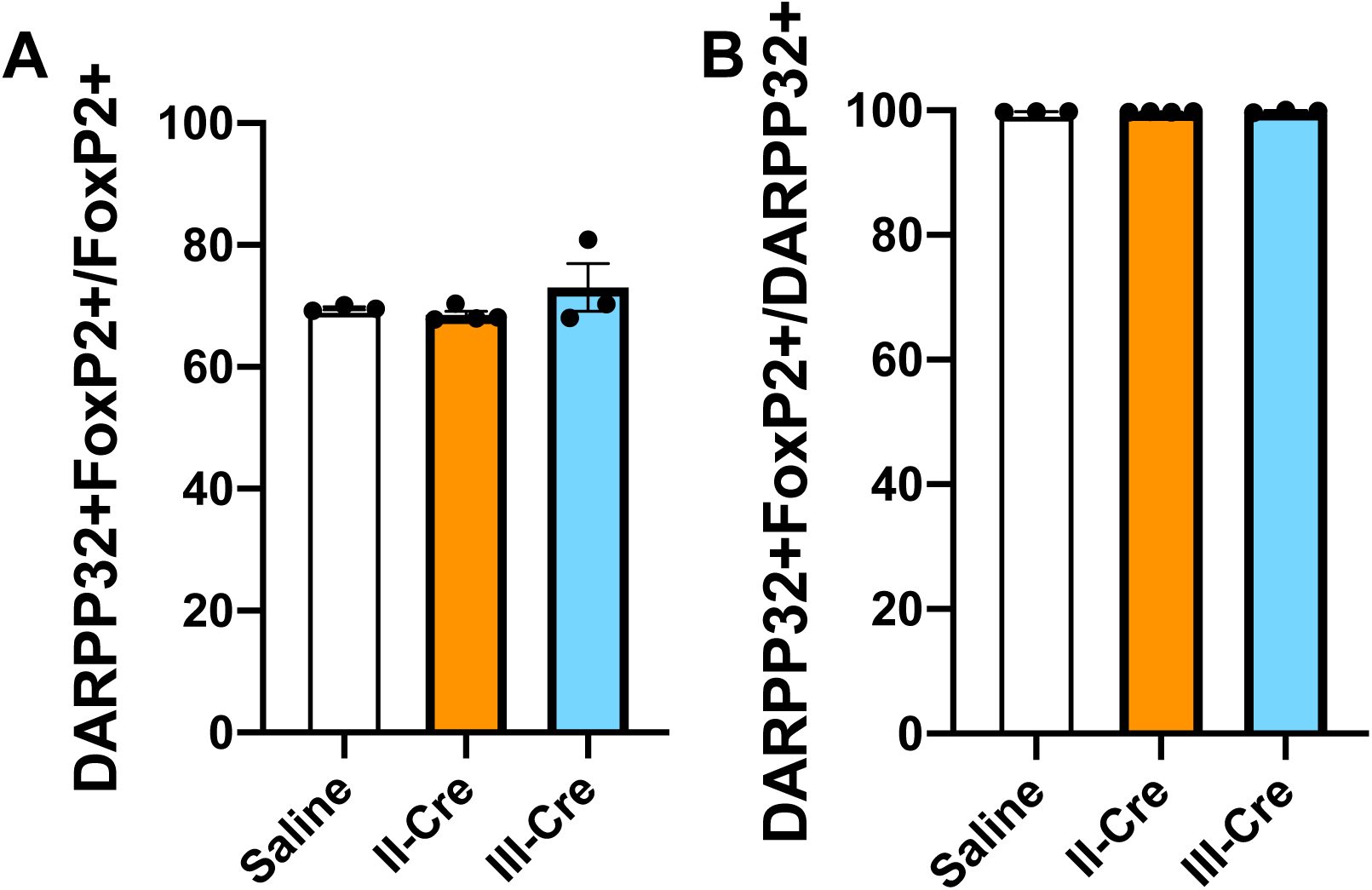
Baseline quantification for DARPP32^+^ and FoxP2^+^ co-localization in the striatum. Brain sections from 3 wpi saline, II-Cre, or III-Cre inoculated mice were labeled for both FoxP2 and DARPP32. (**A)** Percentage of FoxP2^+^ neurons that co-localize with DARPP32 labeling. (**B**) Percentage of DARPP32^+^ neurons that co-localize with FoxP2 labeling. bars = mean ± SEM. N = 9 sections/mouse, 31-44 images/mouse, 3-4 mice/group No significant differences were identified between groups, one-way ANOVA with Dunnett’s post-test.

**Supplemental Figure 5.**
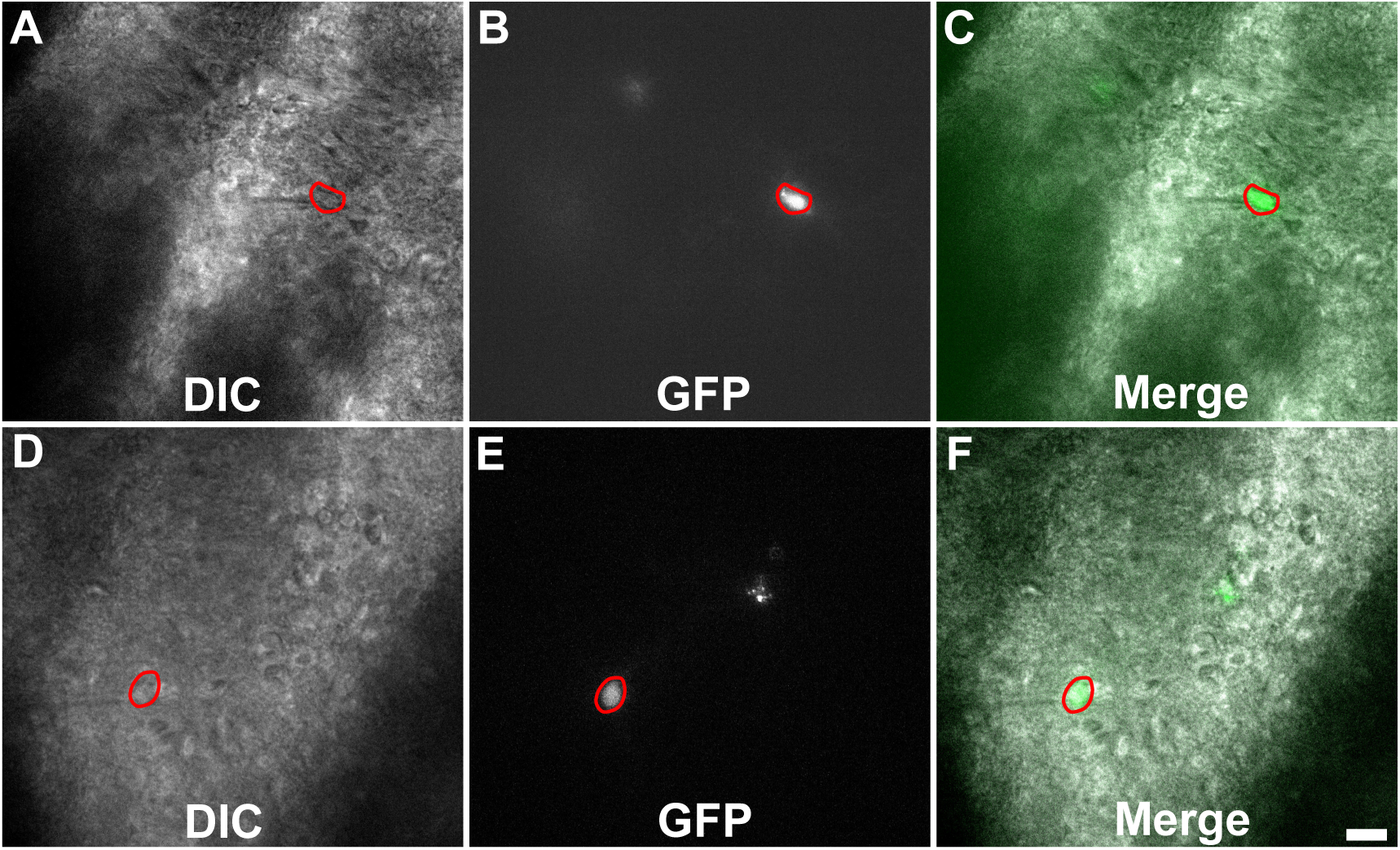
Representative images of TINs used for recording. Images are from 200 *μ*m thick brain sections from III-Cre infected mice that are 10-14 days post infection. Images were obtained on an Olympus BX51. Red circles outline TINs that were patched. Scale bar = 20 *μ*m

**Supplemental Table 1.**
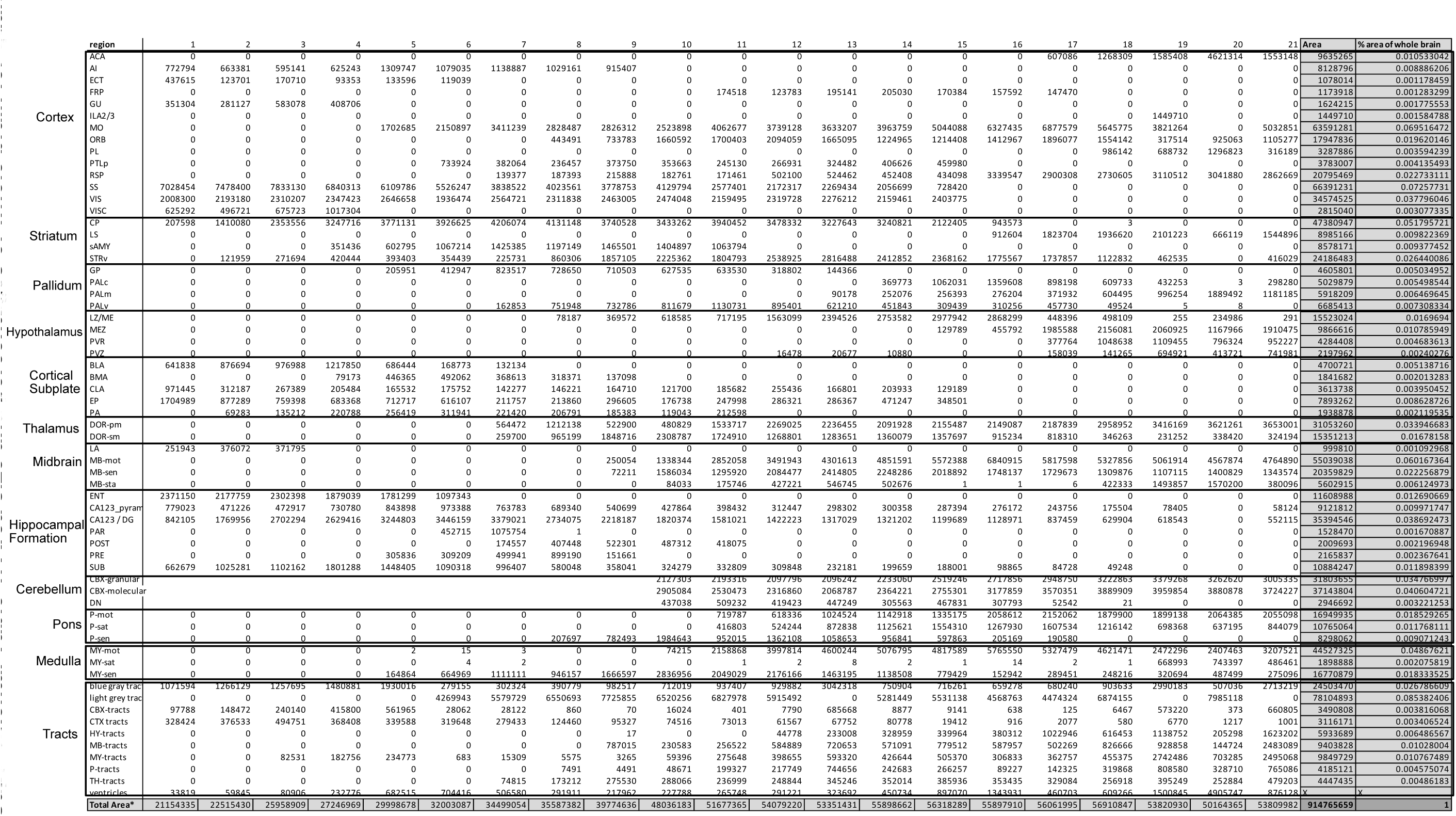
Area size and percentages of whole brain for quantified regions.

**Supplemental Table 2.**
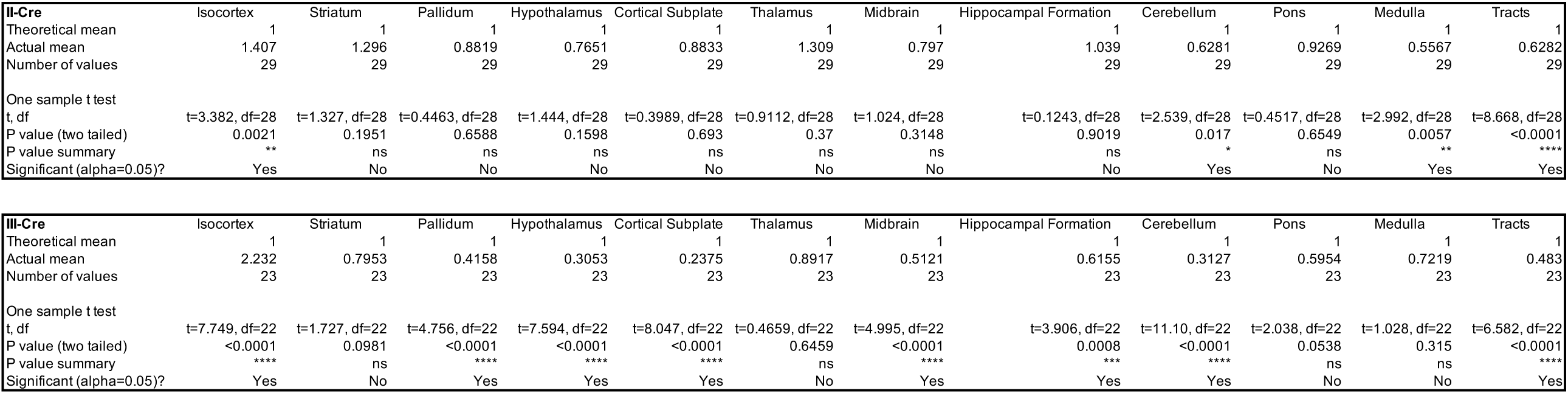
P-values for enrichment scores for the 12 quantified regions.

